# DeepBound: Accurate Identification of Transcript Boundaries via Deep Convolutional Neural Fields

**DOI:** 10.1101/125229

**Authors:** Mingfu Shao, Jianzhu Ma, Sheng Wang

## Abstract

**Motivation:** Reconstructing the full-length expressed transcripts (*a.k.a*. the transcript assembly problem) from the short sequencing reads produced by RNA-seq protocol plays a central role in identifying novel genes and transcripts as well as in studying gene expressions and gene functions. A crucial step in transcript assembly is to accurately determine the splicing junctions and boundaries of the expressed transcripts from the reads alignment. In contrast to the splicing junctions that can be efficiently detected from spliced reads, the problem of identifying boundaries remains open and challenging, due to the fact that the signal related to boundaries is noisy and weak.

**Results:** We present DeepBound, an effective approach to identify boundaries of expressed transcripts from RNA-seq reads alignment. In its core DeepBound employs deep convolutional neural fields to learn the hidden distributions and patterns of boundaries. To accurately model the transition probabilities and to solve the label-imbalance problem, we novelly incorporate the AUC (area under the curve) score into the optimizing objective function. To address the issue that deep probabilistic graphical models requires large number of labeled training samples, we propose to use simulated RNA-seq datasets to train our model. Through extensive experimental studies on both simulation datasets of two species and biological datasets, we show that DeepBound consistently and significantly outperforms the two existing methods.

**Availability:** DeepBound is freely available at https://github.com/realbigws/DeepBound.

## 1 Introduction

RNA-sequencing (RNA-seq) is a well-established and widely-used technology that enables sensitive identification of novel transcripts and accurate measurement of gene expressions (Wang *et al*., 2009). Current high-throughput RNA-seq protocols usually produce short sequencing reads of the expressed transcripts in a given sample. Therefore, a fundamental computational problem is to reconstruct the full-length expressed transcripts from such short sequencing reads, which is usually referred to as the *transcript assembly* problem. Transcript assembly is very challenging, not only because of the difficulties shared with genome assembly problem, such as sequencing errors, homologous repeats, and coverage variations, but more importantly, due to the existence of alternative splicing (in eukaryotes), which drastically increase the complexity of transcript assembly.

Existing assembly methods are either *reference-based* (for example, Cufflinks (Trapnell *et al*., 2010), Scripture (Guttman *et al*., 2010), IsoLasso (Li *et al*., 2011b), SLIDE (Li *et al*., 2011a), CLIIQ (Lin *et al*., 2012), CEM (Li and Jiang, 2012), MITIE (Behr *et al*., 2013), Traph (Tomescu *et al*., 2013), StringTie (Pertea *et al*., 2015), and TransComb (Liu *et al*., 2016b)), or *de novo* (for example, TransABySS (Robertson *et al*., 2010), Rnnotator (Martin *et al*., 2010), Trinity (Grabherr *et al*., 2011), SOAPdenovo-Trans (Xie *et al*., 2014), Velvet (Zerbino and Birney, 2008), Oases (Schulz *et al*., 2012), IDBA-Tran (Peng *et al*., 2013), and BinPacker (Liu *et al*., 2016a)), depending on whether a reference genome is assumed available and being used. Reference-based methods usually give higher accuracy comparing to *de novo* methods. In contrast to the *de novo* methods that gives the nucleotide sequences of the expressed transcripts, reference based methods infer the structures of expressed transcripts, i.e., the coordinates of the *splicing junctions* and *boundaries* (i.e., 5’ and 3’ ends) *w.r.t*. the reference genome for each expressed transcript. To achieve this, reference-based methods first align all the reads to the reference genome using RNA-seq aligners (for example, TopHat2 (Kim *et al*., 2013), STAR (Dobin *et al*., 2013), and HISAT (Kim *et al*., 2015)). Then the reads are clustered into different gene loci based on the alignment, and the coordinates of splicing junctions and boundaries of all expressed isoforms are inferred. Finally, within each gene loci, these coordinates are organized in a so-called *splice graph*, and the expressed transcripts are assembled by computing a set of paths so as to mostly fit the splice graph.

Hence, accurate identification of the splicing junctions and boundaries from the reads alignment is crucial, since they provide building-blocks for transcript assembly. The signal for splicing junctions usually can be clearly reflected in the reads alignment, making the identification of splicing junctions a relatively easy task. Specifically, a splicing junction can be inferred by a group of reads (called *spliced reads*) which are all separately aligned to the reference genome with identical separating coordinates (see Figure 1(a)). In contrast, such strong signal does not exist for transcript boundaries. The intuition that can be used to identify boundaries is that the read coverage tends to increase (resp. decrease) when a transcript starts (resp. terminates). However, such signal is noisy and weak, making the task of identification of boundaries very challenging.

Limited efforts have been focused on inferring boundaries of expressed transcripts from RNA-seq reads alignment. Current reference-based transcript assemblers usually use a simple rule to identify boundaries: the coordinates where reads appear and disappear are identified as start and end boundaries, respectively (hereinafter we call it as TypicalRule). TypicalRule suffers from both high false-positive rate and false-negative rate for the following reasons (Figure 1(**b,c,d**)). First, some boundaries are inside exons, and for such cases TypicalRule is impossible to identify them (Figure 1**(b)**). Second, a region within an exon might not be covered by any reads due to low sequencing depth, high coverage variation, and alignment errors; such gap shall result in TypicalRule reporting two falsepositive boundaries (Figure 1**(c)**). Third, TypicalRule shall also miss two boundaries if contaminations of DNA sequences or misalignments that bridge two transcripts (Figure 1**(d)**). A recent study on meta-assembly employs a more sophisticated method to infer boundaries of the expressed transcripts (Niknafs *et al*., 2016). This method employs Mann-Whitney U test and fold change comparison to identify significant coverage change (hereinafter we call it MWUTest). However, as we will show in this paper, the accuracy of MWUTest is still very low, due to its limited capability to model transitions and to handle noise.

**Fig. 1.**
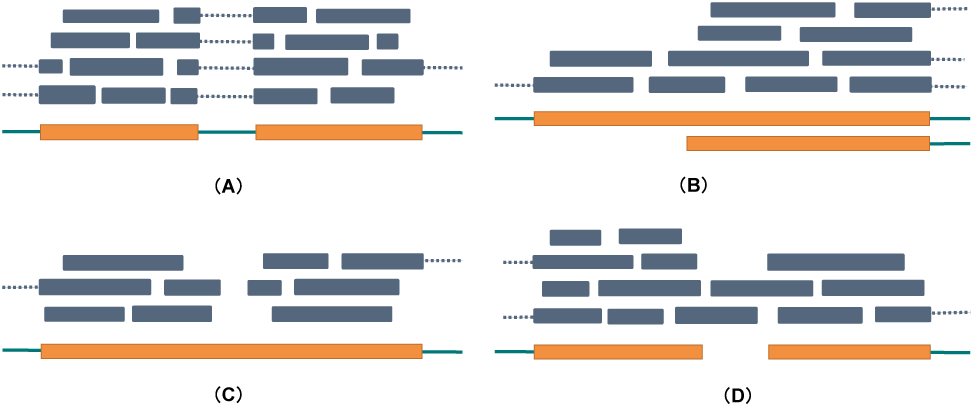
Illustration of identification of splicing junctions and challenging of identifying boundaries. Exons and introns are represented as thick orange bars and thin green bars, respectively. Reads are represented as blue bars, where spliced reads are connected with dotted lines. **(a)** Splicing junction can be inferred from a group of aligned spliced reads. **(b)** A transcript starts in the middle of an exon. In this case we can observe an increasing of read coverage. TypicalRule shall fail to identify this boundary. **(c)** A gap appears in the middle of an exon. TypicalRule shall report two false-positive boundaries. **(d)** Two transcripts are bridges by reads. TypicalRule shall miss these two boundaries.

In this work, we propose a novel algorithmic framework, DeepBound, to identify boundaries of expressed transcripts from reads alignment by using deep convolutional neural fields (DeepCNF) (Wang *et al*., 2015, 2016c). Different from previous statistical methods, DeepBound can integrate a variety of information and automatically determine their quantity under different circumstances. Our model and algorithm are particularly designed so as to take advantage of the properties and to address the challengings of boundary detection. Specifically, first, DeepBound models the boundary detection problem as a sequential labeling problem using a conditional probabilistic graphical model (Lafferty *et al*., 2001). The key idea is to quantify the probability of observing the boundaries on position *i* given the features and states around *i*. Second, given billion level of potential boundaries, patterns of boundaries are complicated. Previous sequential graphical model such as Conditional Random Field and Hidden Markov Model cannot model such complicated scenarios. In contrast, DeepCNF can naturally model complex pattern hidden in the boundaries. Third, as the boundaries are extremely sparse comparing to non-boundary coordinates, previous machine learning models will all be biased towards predicting the input samples as non-boundaries. The inherent reason is that it is very hard to discriminate the true signal with the outliers for sparse observations. To handle this highly label-imbalanced problem, we introduce a novel objective function which directly models the AUC score between the predicted and actual labels (Wang *et al*., 2016a,b). The price we pay is that we get a new objective function which is neither smooth nor convex. However, by applying the Chebyshev approximation (Calders and Jaroszewicz, 2007) on this objective function, we can still be able to optimize it efficiently.

To address the issue that DeepCNF requires large volume of labeled training samples, we novelly propose to use simulation RNA-seq data to train our model. Ideally, realistic parametric distribution of boundaries can be learned more accurately by such a supervised learning way. To valid our assumption and also to compare the performance of our method with that of TypicalRule and MWUTest, we test these methods on simulation datasets from both human and mouse species, as well as on biological RNA-seq datasets. The results demonstrate that the model trained with simulation datasets on one species can also perform well on another species and on biological datasets. On all testing datasets, our methods significantly outperforms other two methods.

## 2 Experimental Results

In this section, we first introduce the experimental pipeline and the evaluation measurements in Section 2.1. After that we then illustrate the performance of all three methods on simulation datasets in Section 2.2, and that on biological datasets in Section 2.3.

### 2.1 Experimental Settings

To identify boundaries of the expressed transcripts in a given RNA-seq dataset, we first align all the reads to the reference genome using the recent RNA-seq aligner HISAT2 (Kim *et al*., 2015). We then cluster reads into different *gene loci* according to their aligned coordinates such that the distance between adjacent gene loci is at least 50bp. Within each gene loci, splicing junctions are identified, defined as two or more spliced reads that share the same splicing coordinates. We use all these splicing junctions as well as the start and end boundaries of the gene loci to partition the whole gene loci into separate *partial exons* (see Figure 2). Each partial exon serves as an independent instance, i.e., each method will take a partial exon (the features of this partial exon, such as coverage profile, sequence information, etc) as input, and identify boundaries of the expressed transcripts within this partial exon. For each dataset and each method, the predicted boundaries by this method for this dataset shall be the union of the identified boundaries for all partial exons in this dataset.

**Fig. 2.**
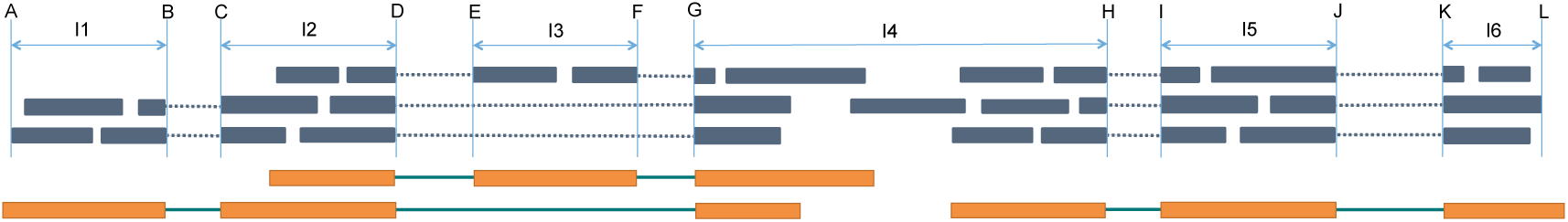
Illustration of partial exons within a gene loci. Thick orange bars (representing exons) and thin green bars (representing introns) show the (unknown) structures of the expressed transcripts. Green bars represent the observed reads that aligned to the reference genome. Notice that the three expressed transcripts come from two genes, while we identify all the reads as a single gene loci (because the single read in the middle of *I*_4_ bridges the two genes). Six splice junctions, (*B, C*), (*D, E*), (*F, G*), (*D, G*), (*H, I*) and (*J, K*), are identified, which, together with the gene loci boundaries, *A* and *L*, partition this gene loci into 6 partial exons, *I_k_*, 1 ≤ *k* ≤ 6. Each partial exon becomes an independent instance. Notice that in partial exon *I*_3_, there exists a gap less than 50bp, thus the two parts are regarded within a single gene loci.

We apply three methods to identify boundaries, TypicalRule, MWUTest, and DeepBound (see Section 3 for details about them). To measure the accuracy of these methods, we first define the ground-truth boundaries. For simulation datasets, we know exactly the expressed transcripts from the simulation process, and their boundaries are served as the ground-truth. For biological datasets, we use the boundaries of all known transcripts in the annotation database, and keep those within a distance of 50bp for some partial exon (in the studied RNA-seq dataset) as ground-truth. Notice that usually for each RNA-seq dataset, only a subset of them gets expressed, and it could be also that some novel transcripts are expressed but not in the current annotation database. Nevertheless, this way of comparison is still fare enough and commonly used to illustrate the comparative performance of different methods.

Given the ground-truth boundaries, we use precision and recall to measure the identified boundaries by each method. We evaluate start boundaries and end boundaries separately. For each dataset, we compute a one-to-one matching between the predicted start (resp. end) boundaries and the ground-truth start (resp. end) boundaries, such that each pair of matched boundaries are adjacent according to their coordinates. We say an identified boundary is *correct* if it is matched to some ground-truth boundary and the distance between them is less than 50bp. With this definition, we can then compute *precision*, which is defined as the ratio between the number of correct boundaries and the total number of identified boundaries, and *recall*, which is defined as the ratio between the number of correct boundaries and the number of ground-truth boundaries. Notice that for our boundary prediction problem it is not appropriate to use true positive and false positive rates. This is because most genomic positions are not boundaries, making the false positive rate always a tiny number. Thus, it is much more meaningful to use precision and recall to perform evaluation.

### 2.2 Results on Simulation Datasets

We use Flux-Simulator (Griebel *et al*., 2012) to simulate the RNA-seq datasets. Flux-Simulator utilizes the known transcriptome annotation database (in our experiments, we use ENSEMBL release-87 annotation) and reference genome to simulate the expressed transcripts and the sequenced reads, following certain empirical distributions. To compare the performance of methods on datasets with different sequencing depths, we use Flux-Simulation to generate two types of datasets, one containing 150M paired-end reads of length 75bp, while the other containing 15M 75bp paired-end reads. To test the methods on different species, we choose two model species, human and mouse, and for each species, we independently simulate 10 datasets for each type of dataset. We independently simulate another two human datasets (with 150M reads in each dataset), and use them to train our DeepCNF model, as well as to determine the parameters in TypicalRule and MWUTest.

The comparison of the three methods on 10 human simulation datasets with 150M reads is illustrated in Figure 3. First, we can observe that DeepBound significantly outperforms both TypicalRule and MWUTest in terms of both precision and recall. Specifically, DeepBound obtains precision of 44% and 59% for start and end boundaries, while TypicalRule (better than MWUTest) gets 27% and 41%, respectively. The advantage of DeepBound is even more pronounced in terms of recall: DeepBound obtains 87% and 81%, while TypicalRule (better than MWUTest) gets 42% and 61%, for start and end boundaries, respectively. Second, MWUTest is better than TypicalRule in terms of recall (about 8.1% and 5.9%, for start and end boundaries, respectively), but slightly worse in terms of precision (about 0.93% and 2.4%). Third, notice that all three methods have a small variation over the 10 datasets, indicating the consistency of the methods on simulation datasets. Fourth, all the three methods behave quite divergently on start and end boundaries. This is probably because the sequencing protocols have different distributions and patterns on start and end boundaries.

The comparison of the three methods on 10 human simulation datasets with 15M reads is illustrated in Figure 4. Again, DeepBound significantly outperforms both TypicalRule and MWUTest, especially in terms of recall. Comparing with the datasets with higher sequencing depth (Figure 3), we can see that precision of all three methods get improved, while recall keeps more or less the same. This is probably because higher sequencing depth also introduces more noise for identifying boundaries. Notice that our model is trained on datasets with 150M reads, thus the results here demonstrate that our method does not suffer from overfitting with the change of sequencing depths.

The comparison of the three methods on 10 mouse datasets are illustrated in Figure 5 and Figure 6 for 150M datasets and 15M datasets, respectively. We can observe that our method trained on human datasets works surprisingly well on mouse, again outperforms TypicalRule and MWUTest by a significant margin. The fact that human-trained model still performs well for mouse tells that the statistical pattern of boundaries is more heavily related with sequencing and alignment errors, and reads coverage bias, rather than different species (notice that although the sequencing reads are simulated by Flux-Simulator, the reads alignment are generated by “real” aligner, HISAT). This is crucial for transcript boundary detection, since, as we illustrate here, we can use deep learning model to capture the hidden patterns related with boundaries from human data, and the trained model shall be general enough to be applied to infer boundaries for other species.

**Fig. 3.**
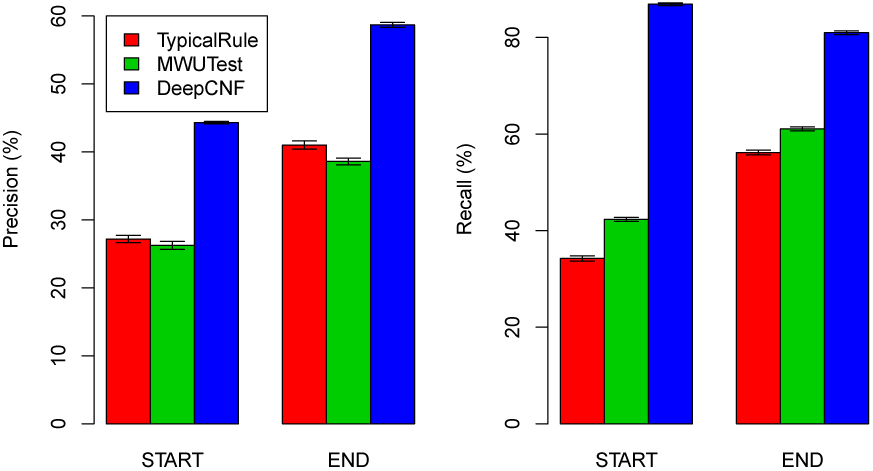
Comparison of the 3 methods on human simulations with each dataset containing 150M reads. The height of each bar shows the average precision (left part) and recall (right part) over the 10 datasets, while the thin bar on top illustrates the standard deviation.

**Fig. 4.**
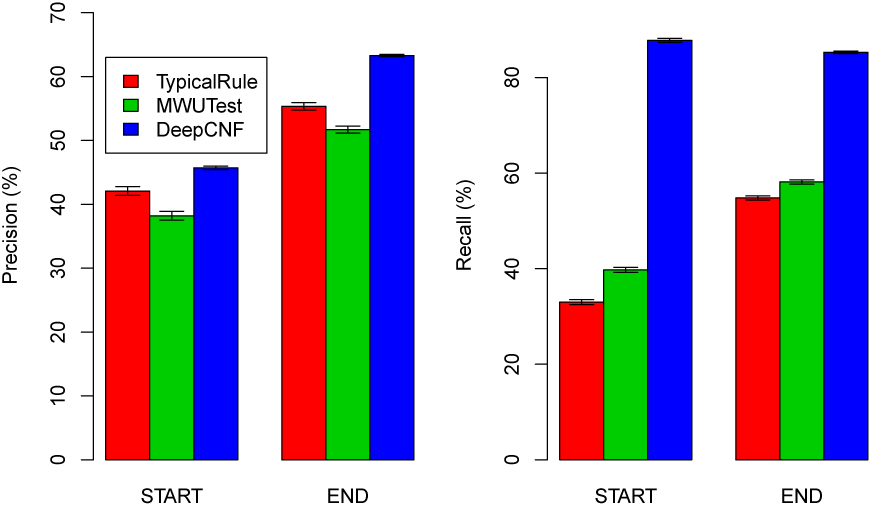
Comparison of the 3 methods on human simulations with each dataset containing 15M reads. The height of each bar shows the average precision (left part) and recall (right part) over the 10 datasets, while the thin bar on top illustrates the standard deviation.

**Fig. 5.**
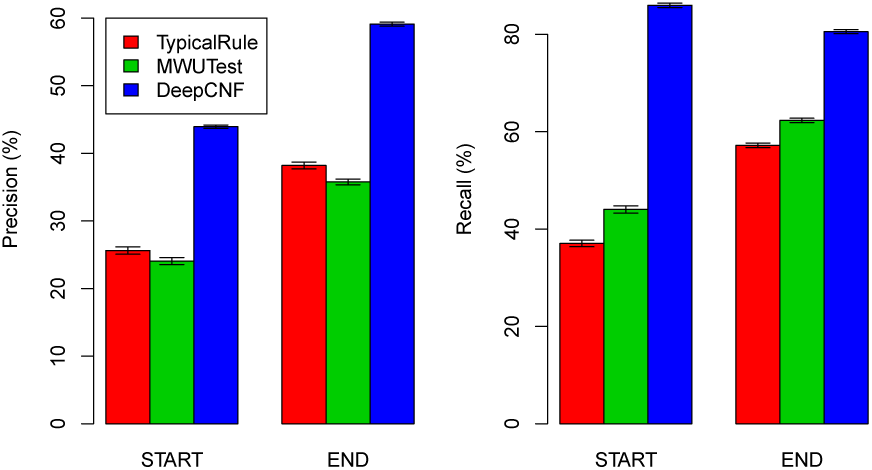
Comparison of the 3 methods on mouse simulations with each dataset containing 150M reads. The height of each bar shows the average precision (left part) and recall (right part) over the 10 datasets, while the thin bar on top illustrates the standard deviation.

## 2.3 Results on Biological Datasets

To compare the performance of the three methods on biological datasets, especially to illustrate whether DeepBound trained on simulation datasets can still obtain high accuracy on biological datasets, we choose 10 RNA-seq datasets from ENCODE project (summarized in Table 1). All these 10 datasets are from human species and employ paired-end and strand-specific sequencing protocols. They have various sequencing depths, read lengths, and come from divergent cell lines.

**Table 1.** Summary of the 10 biological datasets.

| SRA Acc. | GEO Acc. | #Spots | Cell Line | Localization | Length |
| --- | --- | --- | --- | --- | --- |
| SRR534319 | GSM981256 | 25M | CD20+ | cell | 76 |
| SRR534291 | GSM981244 | 114M | IMR90 | cytosol | 101 |
| SRR545695 | GSM984609 | 40M | CD14+ | cell | 76 |
| SRR387661 | GSM840137 | 125M | K562 | cytosol | 76 |
| SRR307911 | GSM758566 | 41M | H1-hESC | cell | 76 |
| SRR545723 | GSM984621 | 147M | HMEpC | cell | 101 |
| SRR315323 | GSM765399 | 30M | NHEK | nucleus | 76 |
| SRR307903 | GSM758562 | 36M | BJ | cell | 76 |
| SRR315334 | GSM765404 | 39M | HeLa-S3 | cytosol | 76 |
| SRR534307 | GSM981252 | 167M | MCF-7 | cytosol | 101 |

The comparison of the three methods on these biological datasets are illustrated in Figure 7. Again, our method outperforms TypicalRule and MWUTest by a significant margin in terms of both recall and precision for both start and end boundaries. These results also demonstrate that our model trained on simulation datasets are not overfitted towards simulations, but still be capable of applying to biological datasets.

**Fig. 6.**
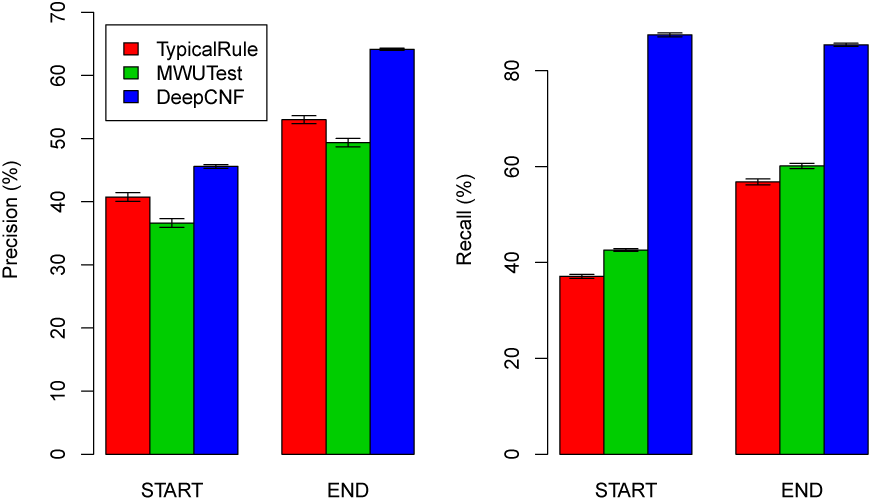
Comparison of the 3 methods on mouse simulations with each dataset containing 15M reads. The height of each bar shows the average precision (left part) and recall (right part) over the 10 datasets, while the thin bar on top illustrates the standard deviation.

**Fig. 7.**
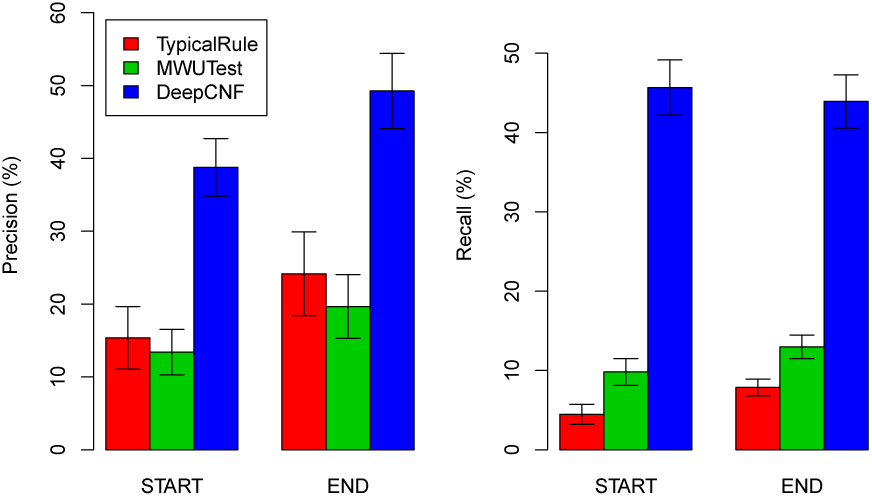
Comparison of the 3 methods on biological datasets. The height of each bar shows the average precision (left part) and recall (right part) over the 10 datasets, while the thin bar on top illustrates the standard deviation.

## 3 Methods

In this Section, we fist describe our method, DeepBound, in Section 3.1. We then give further information about the parameters and implementing details of methodsTypicalRule and MWUTest in Section 3.2 and Section 3.3, respectively.

### 3.1 DeepBound

DeepBound involves three major steps (see Figure 8). Given an instance (i.e., a partial exon), DeepBound first computes features for each position within this partial exon based on the coverage profile, and sequence information around the neighborhood of this position. Second, DeepCNF is applied to learn robust logic that can translate these position-specific features to position-specific probabilities, i.e., the probabilities of being a boundary for each position. Third, with these probabilities, we devise an efficient algorithm to infer actual boundaries, through iteratively identify regions that yield high average probables.

**Fig. 8.**
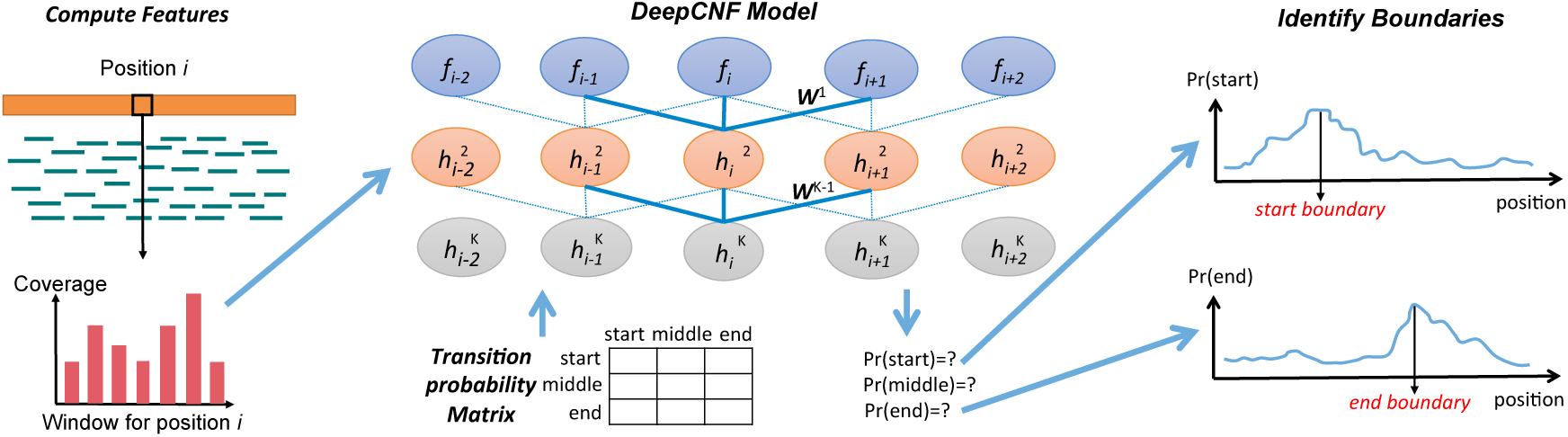
Illustration of the pipeline of DeepBound. Left: Features are extracted from reads alignment. The main feature we are using are the coverage information over a window. Middle: Illustration of the DeepCNF model. DeepCNF model takes all features as input, and predicts the probability of being start, middle, end for each position. Right: The final boundaries are inferred using a greedy algorithm by taking the probabilities from the DeepCNF as input.

#### 3.1.1 Position-Specific Features

For each position *i* within a partial exon we compute 17 features. The first feature we add is the reads coverage for position *i*. We then compute features about the context of position *i* by considering different size of windows around position *i*. Specifically, for a window size of *k*, we compute *n*_*L*_ and *n*_*R*_, which are the number of reads aligned within the region [*i* − *k, i*) and (*i, i* + *k*], respectively. By assuming a null hypothesis of uniform distribution, i.e., each of these (*n*_*L*_ + *n*_*R*_) reads has an equal chance to be in either region, we can compute the pvalue of that left (resp. right) region contains significantly more reads than right (resp. left) region:

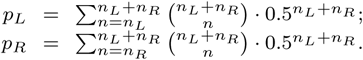

We choose 3 window sizes, *k* = 20, 50, 100, and for each *k*, we add the two pvalues *pL* and *pR* and the two standard deviations of the read coverage in the two regions as features. We also include the sequence information as features. Specifically, for position *k*, we add four binary features, indicating whether the nucleotide at this position is A, C, G, or T. Finally, we incorporate another two binary features, indicating whether position *i* is the start or end boundary of the whole gene loci (the boundaries of gene loci are very likely to be boundaries of expressed transcripts).

#### 3.1.2 AUC-Maximized Deep Convolutional Neural Fields

We formalize the problem of transcript boundary detection as a sequential labeling problem. There are three possible labels, *start, middle, end*, representing whether a position is a start boundary, non-boundary, or end boundary, respectively. For position *i* within the partial exon, its predicted label is determined based on both the features for position *i* and the predicted labels of position (*i* − 1) and (*i* + 1). Our DeepCNF model has two major components: the Conditional Neural Fields (CNF) module (Peng *et al*., 2009), and the Deep Convolutional Neural Network (DCNN) module (Lee *et al*., 2009). The CNF module is a linear chain probabilistic graphical model which explicitly models conditional probability of observing the labelling sequences given the corresponding features. The binary potential of CNF model describes the dependency between the labels of adjacent positions. We substitute the original linear function with the DCNN model to capture the complex patterns in the training data. In particular, given the input features fetched from two adjacent positions, there are nine different DCNN models quantifying the likelihood of observing nine pairs of states (for example, *start* to *middle*, etc), respectively. The weights of all the DCNN models will be learned directly from training data and the nine DCNN models will compete with each other to determine the final assignments for each adjacent positions. Integrating DCNN model with CNF enables us to capture the complicated underlying predicting logic buried in the millions of labeled instances. To control model complexity and to avoid over-fitting, we add an *L*_2_-norm penalty term as the regularization factor and perform 10 fold cross validation to determine the hyper-parameter.

One of the major challenges for the transcript boundary detection is the the number of training samples for different labels are severely imbalanced (Galar *et al*., 2012). Comparing with the non-boundary positions, the boundary positions are much rare, which implies that the empirical estimation of transition probabilities between non-boundary positions and boundary positions will be all nearly zero. This is a common problem for the genome and protein sequence annotation where important nucleotides/genes/residues are rare. Traditional strategies usually solve this problem by down-sampling the training samples with dominating labels. However, in our scenario, down-sampling will break the linear chain model into pieces and lead to a bias estimation of the transition probability. The computational challenge here is to solve the data imbalance problem while still keep the estimation of transition probability unbiased.

To tackle this problem, in this work we replace the widely used log-likelihood objective function of probabilistic graphical model with an estimated AUC objective function during training process (Wang *et al*., 2016a). Note that this objective function is chosen independently to the structure and the potential functions of probabilistic graphical model. AUC can be derived from the ranking of samples by their predicted confidence of different labels (Cortes and Mohri, 2003). The intuition of using AUC objective function is to transform the original classification problem using collection of individual samples to a new pairwise samples ranking problem. That is, instead of trying to assign a particular label *l* to a sample *s*, we try to predict whether sample *s* has a larger marginal probability to be labeled as *l* comparing with sample *t*. It maximizes the AUC score because as long as the partial order of the confidence between any pair of samples maintain the same, the AUC score will be the same. In this application, for a position in a partial exon with label *start*, its predicted marginal probability should be larger than the predicted marginal probability of *middle* and *end* generated by the DeepCNF model. The occurring frequency for each label is then become equal in all the new pairwise ranking samples and hence solve the data imbalance problem.

However, the above AUC-based objective function introduces lots of non-smooth indicator functions in order to indicate the partial order between two samples, making it not concave and thus very hard to optimize. Another obstacle of the AUC function is the pairwise summation which requires *O*(*n*^2^) computing time. To tackle these problems, we first use a 10 order polynomial Chebyshev function to approximate the AUC function as a smooth and differential function (Calders and Jaroszewicz, 2007), which can be computed in linear time. To trade off between performance and training time, here we use the L-BFGS (Liu and Nocedal, 1989) algorithm instead of stochastic gradient descent to optimize it.

We emphasize that the AUC-maximized DeepCNF is a better way to handle data imbalance issue comparing with other methods such as upsampling, down-sampling, or weighting samples. This is because these sampling based methods are designed for independent data points, but not suitable for sequence labeling since performing sampling might break the linear chain structure of the sequence. In fact, we have tried down-sampling on *middle* label positions; the result measured by AUC is only around 0.62, compared to AUC-maximized DeepCNF at 0.93, with the same model architecture. We also performed another experiment, in which we assumed each data is independent and trained a model to maximize likelihood. The resulting AUC was around 0.74, which is still not compatible to 0.93.

We use a three layer neural network with 100 hidden neurons in each layer, and set the window size of features as 75. By trying alternative settings, we found that there is no improvement by using more hidden neurons, more layers, or larger window size regarding the predicting performance. The total 105423 instances (i.e., partial exons) in the 2 human simulation datasets (with 150M paired-end reads in each dataset) are used to train our model.

#### 3.1.3 Predict Boundaries

For each position within a given partial exon, the above DeepCNF model gives the probability of labeling this position as each of the three labels. With these probabilities, we design a greedy algorithm to predict the actual start and end boundaries. Our algorithm iteratively compute the region of length 10bp, which has the largest average probability of label *start* (resp. *end*) boundary. If this average probability is larger than a threshold *p* (we use 10 fold cross validation to fix this parameter as *p* = 0.25), then the middle position of the region is determined as a start (resp. end) boundary. Once a new boundary *i* is determined, the region [*i* − 50, *i* + 50] shall be exempt from identifying the same type of boundary, and the algorithm continues to iteratively process the two sub-partial exons, one from the leftmost position to (*i* − 50), and the other from (*i* + 50) to the rightmost position of the current sub-problem. The algorithm terminates when no new boundary can be found.

### 3.2 TypicalRule

Given a partial exon, TypicalRule first partitions it into sub-partial exons by identifying gaps (i.e., regions with read coverage of 0) of length larger than a threshold *g*. Then for each such gap, TypicalRule reports two boundaries, the left position of the gap as the end boundary, and the right position of the gap as the start boundary. In addition to that, if the leftmost (resp. rightmost) position of the given partial exon is the start (resp. end) boundary of the whole gene loci, then this position shall be reported as a start (resp. end) boundary. We have tried different value of *g* ∈{5 · *k* | 1 ≤ *k* ≤ 9} on the two training human simulation datasets, and choose *g* = 10.

### 3.3 MWUTest

MWUTest is a subroutine used in the TACO package to detect *change point* (see (Niknafs *et al*., 2016) Online Methods), which is essentially the same problem we study here to identify transcript boundaries. Given a partial exon, MWUTest iteratively identifies the *potential* change point, defined as the position *i* that minimizes the following mean square error (MSE):

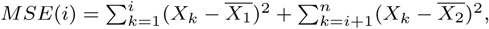
where *X*_*k*_ is the reads coverage at position *k, n* is the total length of this partial exon, and *X*_1_ (resp. *X*_2_) is the average reads coverage for the region [1, *i*] (resp. (*i, n*]). MWUTest then tests whether this potential change point *i* meets both the following two significance criteria: the pvalue under the Mann-Whitney U test shall be less than a threshold (we use the two training datasets to determine this parameter as 0.00001), and fold change of the average coverage between the two sides shall be larger than a threshold (we fix it as 3.0 through trying different values on the training datasets). If the potential change point satisfies these two criteria, then MWUTest shall report this position as a transcript boundary, and iteratively process the two sub-partial exons. The algorithm terminates if no potential change point meets these two criteria.

## 4 Conclusion and Discussion

In this paper, we propose a new approach, DeepBound, to identify boundaries of expressed transcripts from RNA-seq reads alignment by using deep convolutional neural fields. The performance of our method has been extensively evaluated on both simulation datasets and biological datasets, and the results demonstrate that our method significantly outperforms two existing methods for boundary detection.

We emphasize that our model is trained on the reads alignment generated by simulated RNA-seq reads (using Flux-Simulation) followed by aligning them with “real” aligner (HISAT). We show that this training process is successful by illustrating that this model can also achieve high accuracy when applied to other species and to biological datasets. This opens a new way to employ the power of deep learning technologies to solve challenging sequencing problems when large-scale labeled training samples are not available.

Our learning framework is general and can be easily adapted to add other features and trained and applied to other species. We have provided the training and testing software package and users can use their own gene boundary datasets to train their own model. In this work we only apply RNA-seq because it is the most general and popular protocol.

A natural extension of our method is to incorporate it into the complete pipeline of transcript assembly. As a stand-alone software package, DeepBound can be used to detect transcript boundaries in the RNA-seq experiments, and the results can be further applied to correct transcript abundance estimation and to study gene alternative splicing. For example, recent genomic analyses have indicated that many eukaryotic genes can have quite complex alternative splicing products, such as transcript embedded within another transcript (Kumar, 2009), transcripts from reverse strand (Adelman *et al*., 1987), intron retention (Ner-Gaon *et al*., 2004), and polyadenylation sites (Tian and Manley, 2013). We shall apply our tool to analyze these cases in future.

## Acknowledgements

MS was supported in part by the Gordon and Betty Moore Foundation’s Data-Driven Discovery Initiative Grant GBMF4554, the US National Science Foundation Grants CCF-1256087 and CCF-1319998, and the US National Institutes of Health Grant R01HG007104 to Carl Kinsford. SW was supported in part by the US National Institutes of Health Grant R01GM089753 and the US National Science Foundation Grant DBI-1564955 to Jinbo Xu.

